# Distributed source modeling of intracranial stereoelectroencephalographic measurements

**DOI:** 10.1101/2020.08.18.256669

**Authors:** Fa-Hsuan Lin, Hsin-Ju Lee, Jyrki Ahveninen, Iiro P. Jääskeläinen, Hsiang-Yu Yu, Cheng-Chia Lee, Chien-Chen Chou, Wen-Jui Kuo

## Abstract

Intracranial stereoelectroencephalography (sEEG) provides unsurpassed sensitivity and specificity for human neurophysiology. However, sEEG group analyses are complicated because the electrode implantations differ greatly across individuals. Here, using an auditory experiment as the test case, we developed a distributed, anatomically realistic sEEG source-modeling approach for within- and between-subject analyses. In addition to intracranial event-related potentials (iERP), we also estimated the sources of high broadband gamma activity (HBBG), a putative correlate of local neural firing. The source models accounted for a significant portion of the variance of the sEEG measurements in leave-one-out cross-validation. After logarithmic transformations, the sensitivity and signal-to-noise ratio were linearly inversely related to the minimal distance between the brain location and electrode contacts (slope≈-3.6). The HGGB source estimates were remarkably consistent with analyses of intracranial-contact data. In conclusion, distributed sEEG source modeling provides a powerful neuroimaging tool, which facilitates anatomically-normalized group analyses of both iERP and HBBG.

## Introduction

Stereoelectroencephalography (sEEG) detects the neural activity by measuring the electric potentials recorded from electrodes placed inside the brain. Different from electroencephalography (EEG) and magnetoencephalography (MEG), where the neurophysiological signals are measured extracranially, the intracranial recording allows sEEG to provide unprecedented sensitivity to detect local neural currents ^1^. In addition to being used to localize epileptogenic zones ^2,3^, sEEG has also been used for high-level cognitive function studies ^4–6^. While the practical procedure for localizing the electrodes and the associated data analysis has been reported ^7^, a systematic way to combine sEEG data from different patients to summarize effects observed from discrete sampling loci is still lacking. This difficulty is the consequence of optimizing electrode locations based on individual patients’ medical history, imaging data, and vasculature ^8–10^.

One way to address this challenge is to model the spatial distribution of neural currents based on sEEG data in each patient. In EEG and MEG studies, distributed source modeling ^11^ estimates a spatial distribution of neural current consistent with the measurements over a defined “source space”, which is typically over the cortical surfaces, because MEG and EEG have a limited sensitivity beyond the superficial depth ^12^. Individually estimated neural current distributions can be spatially registered to a common coordinate system to derive group-level inferences. This framework has been extensively used in MEG and EEG data analysis ^13^. An early attempt of distributed source modeling of sEEG estimated the origin of the evoked auditory response ^14^. However, the brain was only modeled as a simple one-layer sphere and the analysis was done in individual patients. Simulations based on an infinitely homogeneous model using a solver closely related to the Minimum-Norm Estimates (MNE) suggested that accurate localization of focal neural activity can be achieved if there are electrodes no farther than 15 mm ^15^. However, estimating the source of sEEG data using an anatomically realistic head model and empirical measurements and combining the modeling results across patients remains unexplored. A comprehensive description of the procedure and a quantitative evaluation of the performance will have broad impact in bringing this tool in the neuroimaging community.

One of the most interesting sEEG phenomena, not accessible to scalp EEG, is high-frequency (> 50 Hz) broadband gamma activity (HBBG). In contrast to large-scale neural oscillations visible also to scalp EEG and MEG, HBBG is believed to reflect “non-oscillatory” broadband signals, which not only reveal high-frequency synaptic effects but also provide a direct correlate of local spiking activity ^16–18^. In non-invasive recordings, sensory responses at comparable frequencies are typically much briefer and/or weaker than the sustained HBBG observable in intracranial recordings ^19,20^. Measurements of HBBG could thus offer an opportunity for much more detailed neurophysiological inferences than those based on iERP or oscillatory local field potentials (LFP) alone ^21^. What makes the HBBG specifically interesting from a source modeling perspective is that these signals do not seem to spread far from their neural site of origin. However, no previous study has, to our knowledge, examined HBBG using sEEG source modeling analyses.

Here we report a comprehensive workflow to allow for the distributed modeling of neural currents, including both iERP and HBBG patterns, using sEEG with electrode locations optimized for individual patients with realistic head models. The spatial distributions of neural currents were estimated by the MNE ^22^. Importantly, we quantitatively studied three issues related to the modeling. First, we depicted and calculated spatial distributions of sensitivity and SNR of sEEG. Then, based on empirical sEEG measurements, we used a cross-validation procedure to evaluate how much information can be estimated by the distributed source modeling. Third, considering patient’s safety and the access to imaging resources, different types of data (CT and/or MRI) are used in identifying the locations of electrode contacts. We quantified how the variability in the estimated locations of electrode contacts affect the spatiotemporal characteristics of the estimated neural currents. This distributed source modeling framework and performance assessment are expected to facilitate studies deriving neuroscience inferences using sEEG measurements from patient groups.

## Results

### Locations of electrode contacts

**Figure 1 (A)** shows the locations of electrodes, locations for the neural current estimates, and the head models from a representative patient. The locations of all electrodes from all patients on a standard template (MNI 305) are shown in **Figure 1 (B)**. Electrodes were distributed mostly around the temporal lobes and frontal lobes. Electrodes were implanted in both left and right hemispheres.

**Figure 1.**
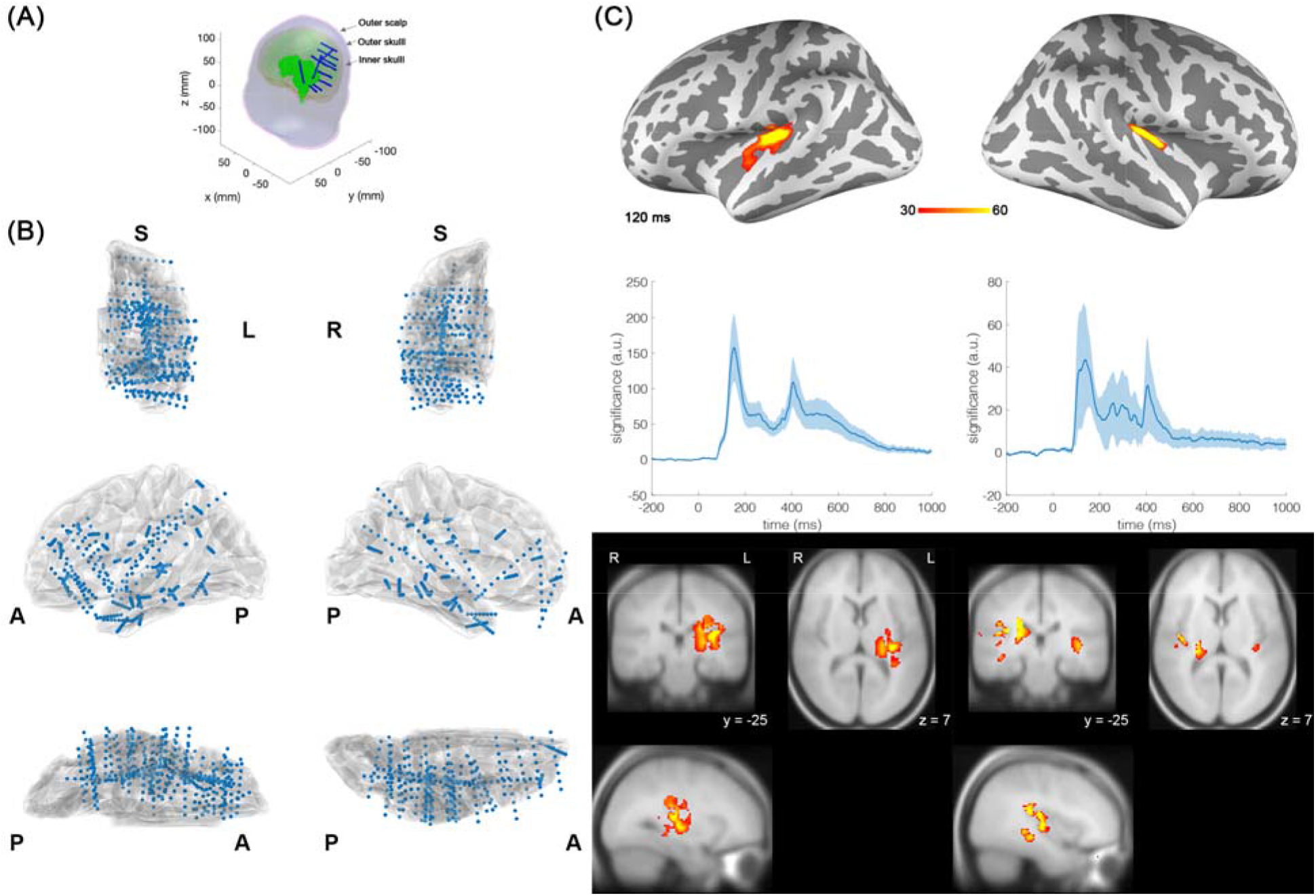
Electrodes, brain models, and the estimated group-averaged neural current distributions and dynamics elicited by the auditory stimulus. (A) An illustration of the source space (green dots) including cortical and sub-cortical brain areas, locations of the electrode contacts (blue dots), and three anatomical boundaries (inner skull, outer skull, and outer scalp) for the lead field calculation using Boundary Element Model on a representative patient. (B) Locations of all electrode contacts (blue dots) depicted over a template brain volume (MNI305) in the left (left column) and right (right column) hemisphere. A: anterior. P: posterior. L: left. R: right. (C) Spatial distributions of significant neural current estimates at 120 ms after the auditory stimulus at the group level. Left and right hemisphere results were selectively derived from patients with electrodes implanted in the left and right hemispheres, respectively. The distributions over an inflated brain model (top row) and at orthogonal anatomical slices (bottom row; L: left hemisphere) are shown at the left and right columns, respectively. The average and the standard deviation of the significance time course within the auditory cortex determined from all patients are shown in the right columns.

### Estimated neural current distributions and dynamics

The estimated neural currents in response to the auditory stimuli from two representative patients are shown in **Figure 2**. Strong activity starting at around 120 ms was found at the left and right temporal lobes for representative patients with electrodes at the left and right temporal lobes, respectively. The spatial distribution and the waveform of the source estimates averaged across patients were shown in **Figure 1 (C)**.

**Figure 2:**
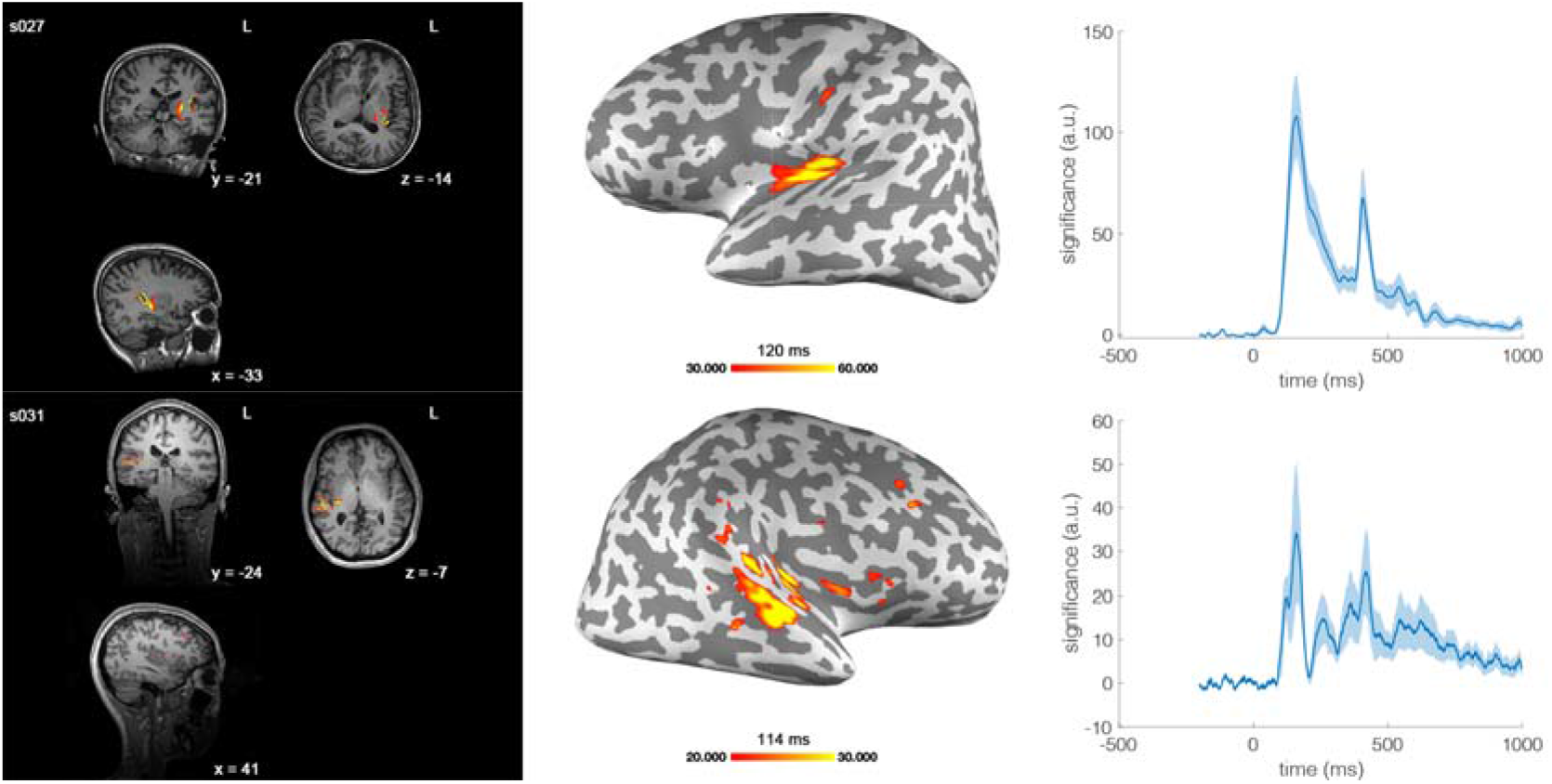
Spatial distributions of significant neural current estimates at about 120 ms after the sound onset in two representative patients with electrodes implanted in left (s027) and right (s031) hemispheres. These distributions at orthogonal anatomical slices (L: left hemisphere) and over an inflated brain model are shown at the left and right columns, respectively. The average and the standard deviation of the significance time course within the auditory cortex determined from all patients are shown at the right columns.

### High-frequency broadband gamma (HBBG) activity

We examined the spatial distribution of HBBG due to the auditory stimulus between 60 Hz and 140 Hz. **Figure 3** shows average spatial distributions of significantly increased and decreased HBBG in STG and IFG from patients with electrodes implanted in the left hemisphere, respectively. These HBBG power changes were further validated from the electrode measurements of three patients, who had electrodes implanted in both left STG and IFG **(Figure 3)**. Similar patterns of significantly increased and decreased HBBG were observed by leaving either one of the three validation patients away from the group average analysis. The source modeling results matched the finding at the electrode around the left STG and IFG by showing significantly increased and decreased HBBG power, respectively.

**Figure 3.**
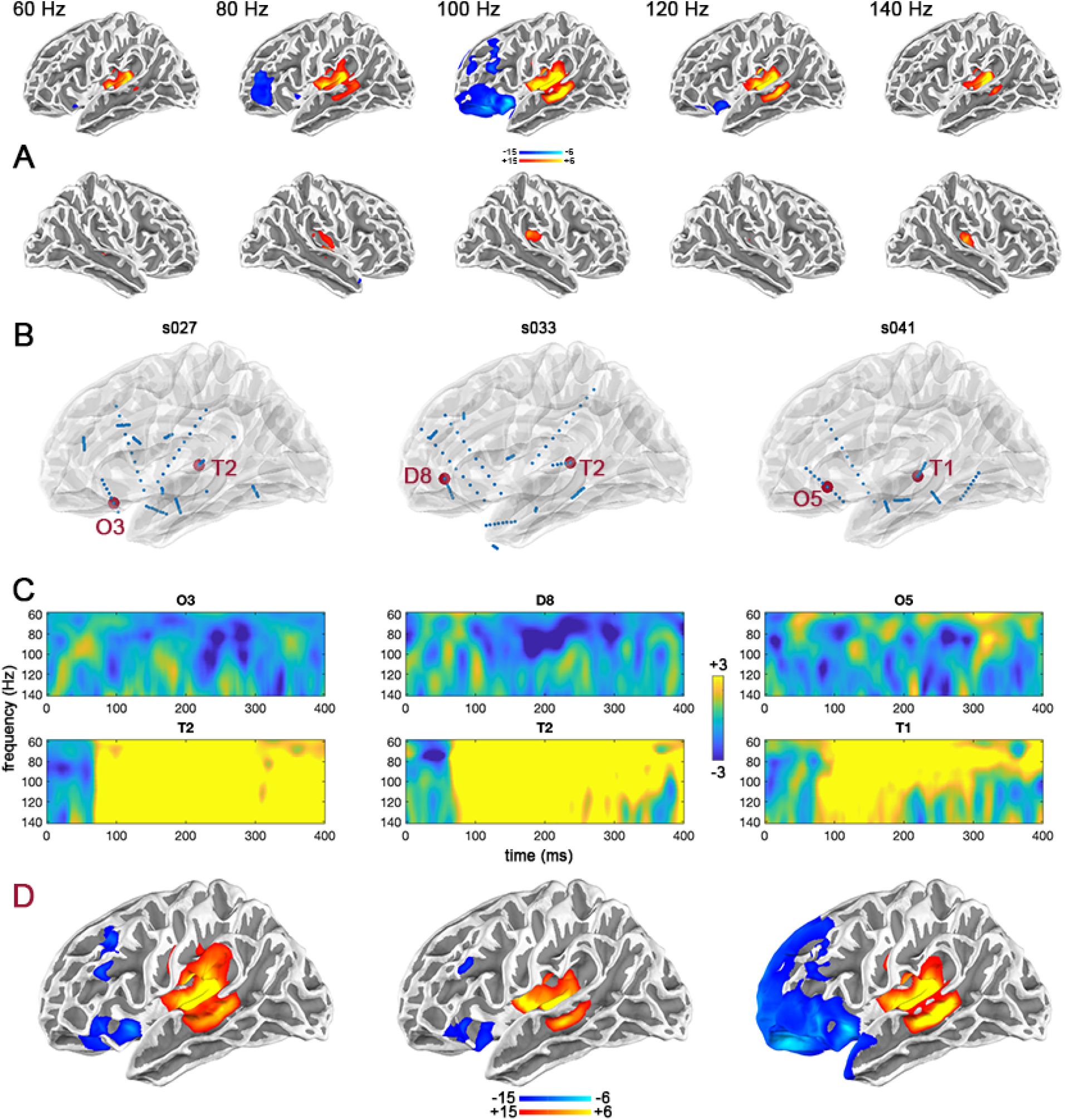
Spatial distributions of HBBG estimates between 60 Hz and 140 Hz in response to the auditory stimulus. (A) Significantly increased auditory HBBG activity were found in STG, whereas the HBBG in IFG was significantly decreased. (B) The recordings from three patients with electrodes implanted at STG and IFG were used to validate the source analysis result: The dark red circles show the locations of the electrode contacts closest to the STG and IFG locations of interest, the blue circles denote the rest of the electrode contacts. (C) The time-frequency representations (TFR) of the data from the electrode T, which is close to STG, show evidence of significantly increased HBBG activity at about 100 Hz between 200 ms and 400 ms after the stimulus onset. The data from the electrodes O and D, which is near IFG, show evidence of significant HBBG decreases, respectively. (D) Leave-one-out between-subject cross-validation analyses: The TFR results from the electrodes T, D, and O were consistent with the source modeling of the estimated HBBG power at the group level. Notably, each of the group representations at the bottom panel include all patients except the one whose electrode data was utilized for the between-subject cross-validation.

### Cross-validation

Distributed source modeling provides estimates of spatially distributed currents. As a model, the estimated neural current is different from the true neural current. Using leave-one-out validation, we quantified this discrepancy **(Table 1**). The average proportion of the explained variance dropped from 38% to 24% as fewer (from 100% to 50%) remaining electrode contacts were used for modeling the source. Depending on the number of electrode contacts used for modeling, the explained variance at an electrode contact ranged between 49% and 22%. No significant relationship between the number of electrode contacts and the proportion of the explained variance was found. This was presumably due to the fact that the number and location of electrodes for each patient were planned based on the clinical need rather than the auditory study.

**Table 1.** Percentages of the variance explained by the source modeling in the cross-validation analysis.

| Patient | Percentage of remaining contacts ($N-1$) for source modeling | | | | Number of electrode contacts ($N$) | Electrode at hemisphere | |
| --- | --- | --- | --- | --- | --- | --- | --- |
|  | 100% | 90% | 70% | 50% |  | left | right |
|  |  |  |  |  |  | hemi. | hemi. |
| s026 | 38% | 34% | 23% | 28% | 69 | + | - |
| s027 | 36% | 32% | 32% | 17% | 104 | + | - |
| s031 | 22% | 21% | 18% | 18% | 95 | - | + |
| s032 | 33% | 31% | 30% | 26% | 67 | + | - |
| s033 | 44% | 45% | 34% | 30% | 76 | + | - |
| s034 | 36% | 33% | 32% | 27% | 100 | - | + |
| s036 | 49% | 37% | 37% | 19% | 59 | - | + |
| s041 | 42% | 34% | 22% | 19% | 70 | + | + |
| max. | 49% | 45% | 37% | 33% | 104 |  |  |
| min. | 22% | 21% | 18% | 17% | 59 |  |  |
| average | 38% | 33% | 29% | 24% | 80 |  |  |
| standard deviation | 8% | 7% | 6% | 7% | 17 |  |  |

Numbers for each patient and the average, maximum, minimum, and standard deviation across patients are reported. The presence and absence of the electrodes at hemispheres were denoted by “+” and “-”, respectively.

### Sensitivity

The spatial distributions of SNR and sensitivity across patients in the left and right hemispheres are shown in **Figures 4** and **5**, respectively. Color coded the brain locations with SNR or sensitivity between the top 85% and 99% of the values across the cortex. Note that only patients with electrodes implanted at one hemisphere were included in the plot of the same hemisphere. Approximate locations of electrode contacts to the cortex were indicated by green dots in the figure. Regions with high SNR and sensitivity were found in the vicinity of electrode contacts. Within an individual, the spatial distributions of SNR and sensitivity were very visually similar, suggesting that the estimated noise distributions were rather spatially homogeneous. The SNR and sensitivity in the thalamus and brain stem were comparable to those locations at the vicinity of electrode contact implantation: they were about the top 85% of the SNR and sensitivity. **Figure 6** shows the spatial distributions of average SNR and sensitivity across patients with electrodes implanted in the left and right hemispheres in brain surfaces and volumes, respectively. Top 15% values were color-coded. High SNR and sensitivity were found around the anterior temporal lobe, insula, and frontal lobe, where electrodes were implanted based on the clinical need to ascertain epileptogenic zones. Both medial and lateral aspects of the cortex and deep brain areas, including part of the brain stem, had high SNR and sensitivity.

**Figure 4.**
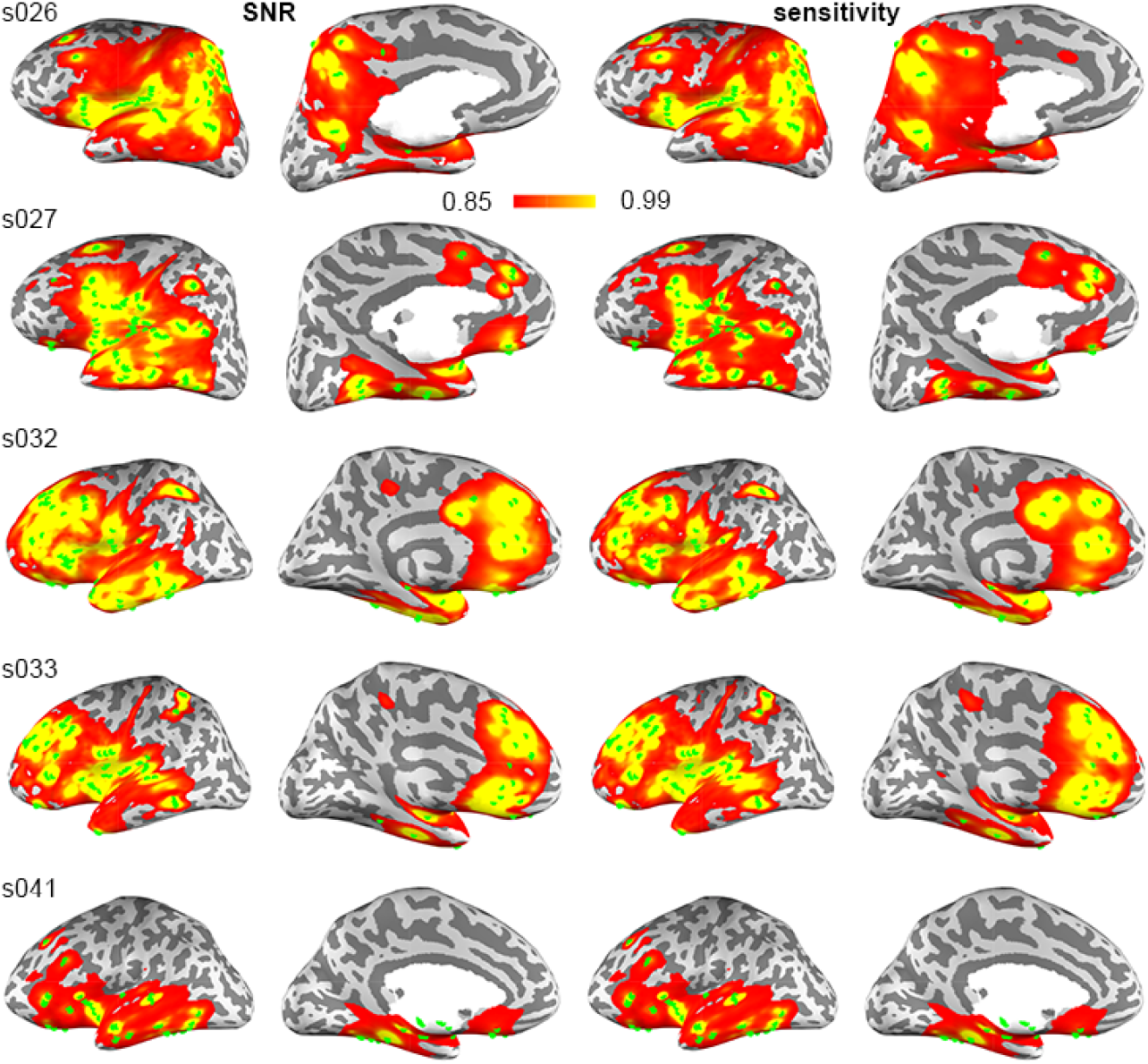
Distributions of the SNR and sensitivity for patients with electrode contacts in the left hemisphere. Colors code values sorted between the top 85% and 99% of each individual patient. Green dots denote the location of electrode contacts.

**Figure 5.**
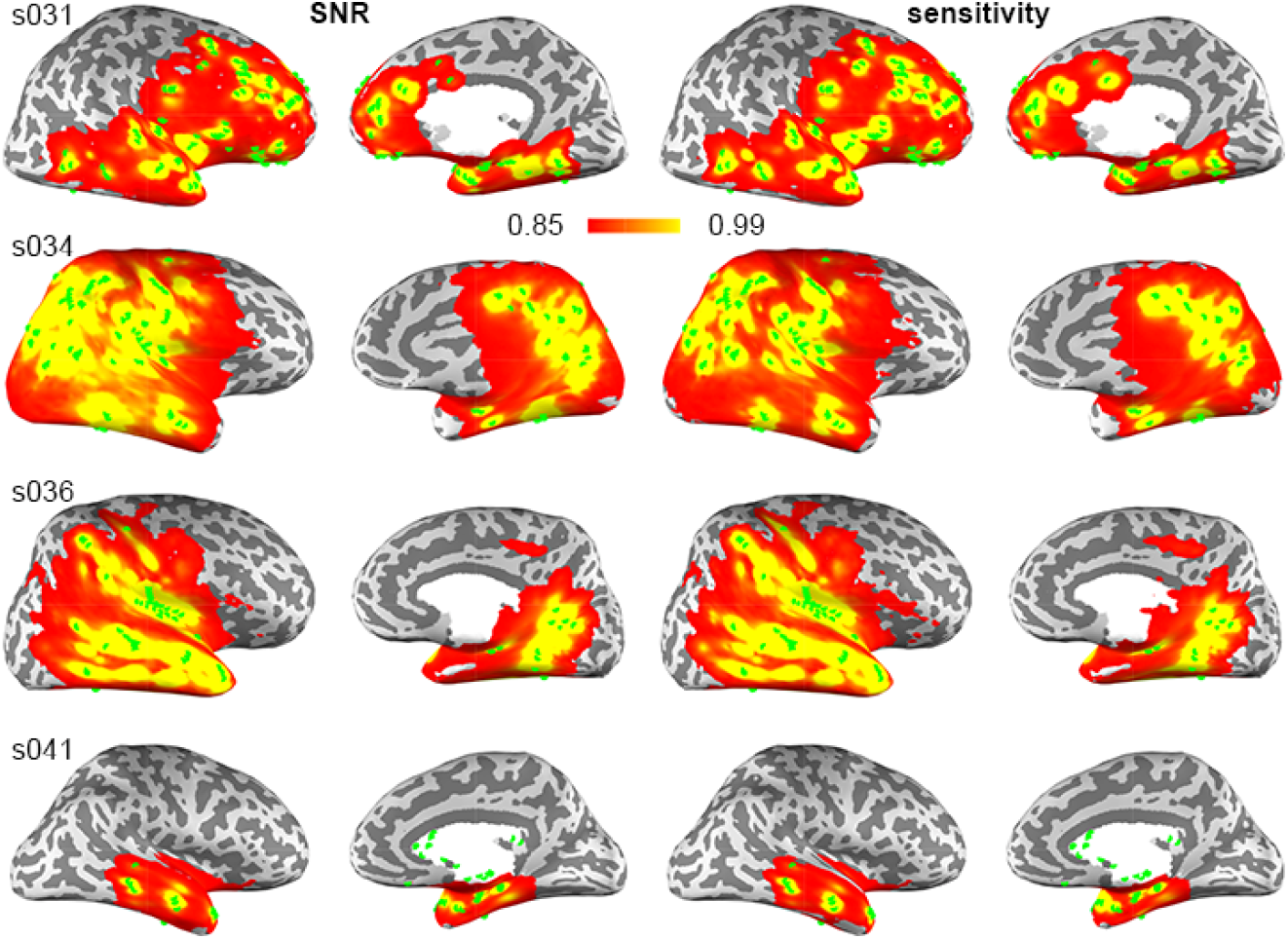
Distributions of the SNR and sensitivity for patients with electrode contacts in the right hemisphere. Colors code values sorted between the top 85% and 99% of each individual patient.

**Figure 6.**
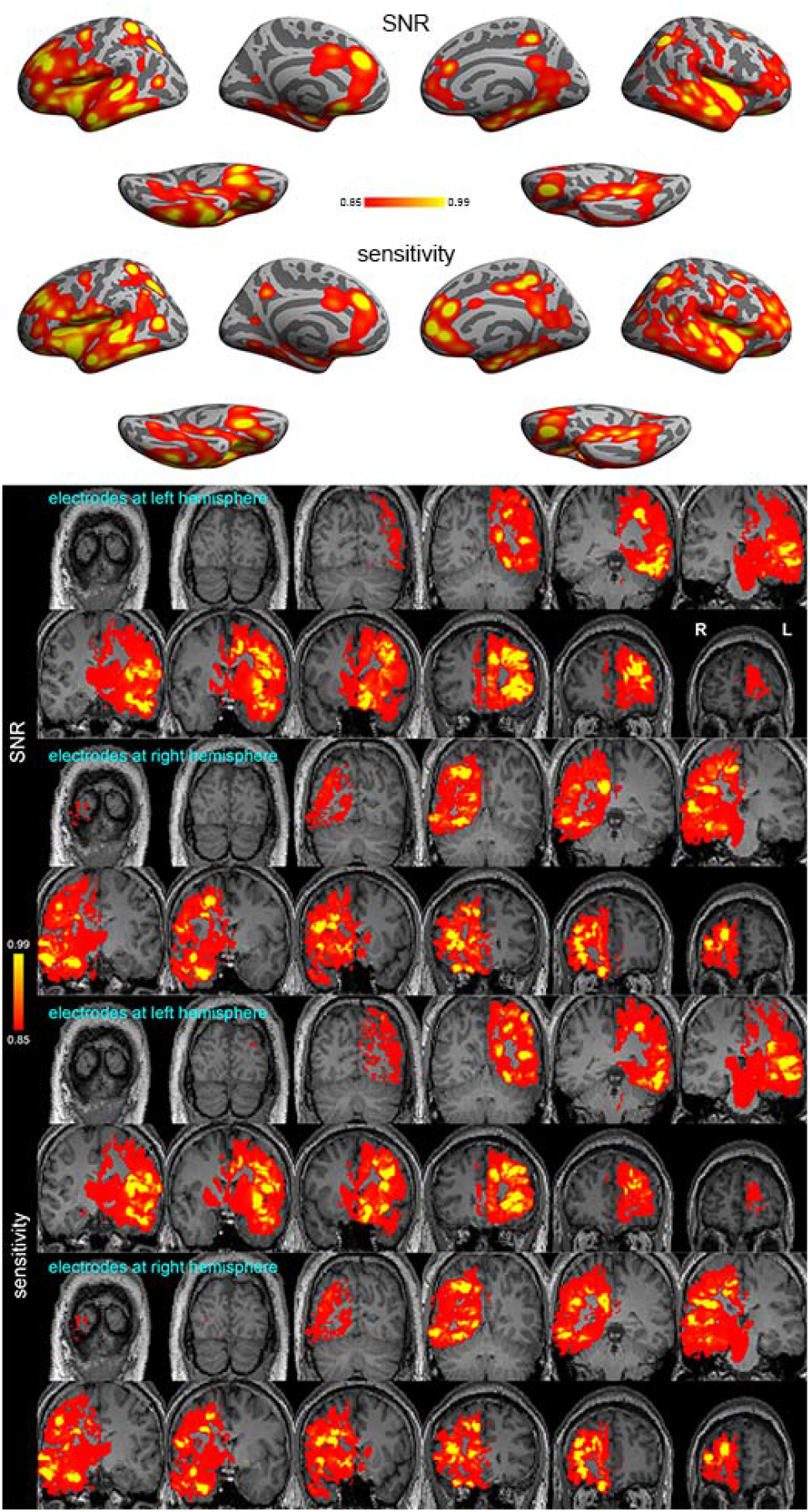
Distributions of the average SNR and sensitivity across patients with electrodes implanted in the left and right hemispheres. Colors code values sorted between the top 85% and 99% of the average.

We quantitatively characterized how the SNR and sensitivity at a brain location varied with its distance to electrode contacts. Specifically, we considered two distance metrics, the minimum and the median of all distances between a brain location and all electrode contacts. The distributions of both distance metrics versus SNR and sensitivity for each patient and all patients in the logarithm scale were shown in **Figures 7** and **8** respectively. These distributions were visually similar across patients.

**Figure 7.**
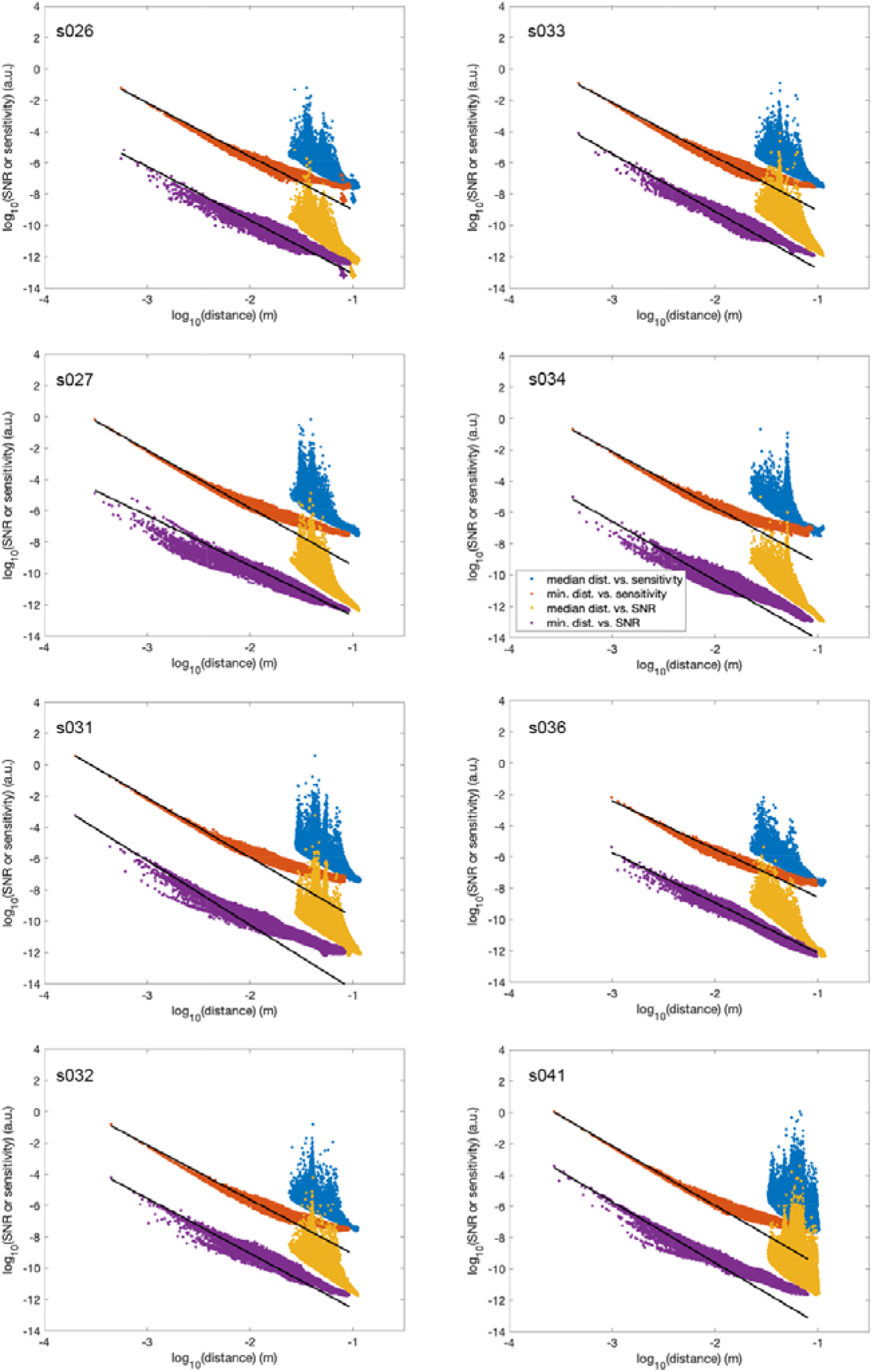
Scatter plots between the brain-electrode distance and SNR and sensitivity in individual patients in the log-log scale. The distance measures include the median and the minimum distances between the estimated brain area and electrode contact locations. Regression lines between the minimum distance and the SNR or sensitivity are also shown.

**Figure 8.**
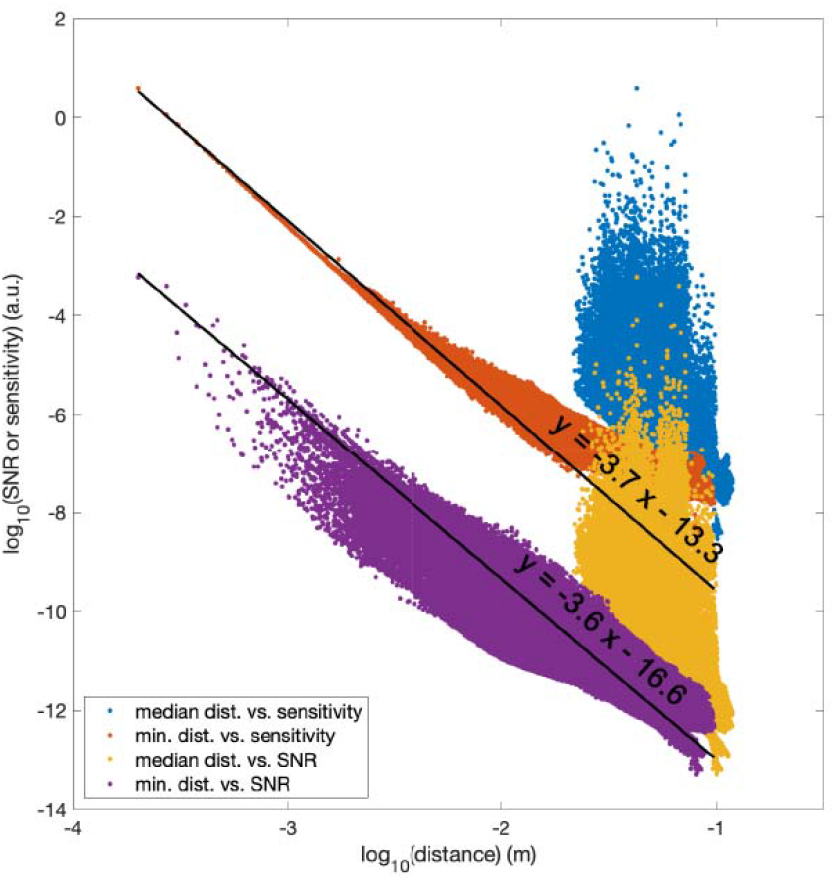
Scatter plots between the brain-electrode distance and SNR and sensitivity across all patients in the log-log scale. The distance measures include the median and the minimum distance between the source-estimated brain area and electrode contact locations. Regression lines between the minimum distance and the SNR or sensitivity were shown.

In the logarithm scale, the distributions of SNR and sensitivity versus the minimum of the distance appeared to be better described by an inverse linear model than the distributions of SNR and sensitivity versus the median of the distance. The regression identified a significant inverse linear relationship between the 10-based logarithm of the SNR and the 10-based logarithm of the minimum distance with a slope of −3.6 and an intercept of −16.6 (*p* < 0.01). A significant inverse linear relationship between the 10-based logarithm of the sensitivity and the 10-based logarithm of the minimum distance was found with a slope of −3.7 and an intercept of −13.3 (*p* < 0.01). Both regression lines suggested that when the minimal distance became tenfold larger (*e.g*., 1 mm to 10 mm), the SNR and sensitivity dropped to 0.25% and 0.20%, respectively.

### The effect of the variability in electrode contact locations

The locations of electrode contacts informed by CT and post-surgery MRI on two representative patients are shown in **Figure 9**. While small, the discrepancy between the estimated locations of electrode contacts by CT and post-surgery MRI was still visually discernable. The average, standard deviation, maximal, and minimal distance between electrode contacts informed by CT and post-surgery MRI were and 1.39 mm ± 0.63 mm, 2.87 mm, and 0.24 mm for one patient and 2.61 ± 0.83 mm, 4.33 mm, and 0.92 mm for the other patient. The dSPMs estimated from two lead field matrices with electrode contact locations informed by CT or post-surgery MRI were shown in **Figure 9**. The spatial distributions and the waveform of the significance of the neural currents were similar. Quantitatively, the correlation coefficients of the spatiotemporal dynamics between two lead field matrices were both 0.97 in two patients. The ratios of the power of the dSPM difference between two lead field matrices and the average power of the dSPM between two lead field matrices were 3.34% and 4.04% for patient s026 and s041, respectively.

**Figure 9.**
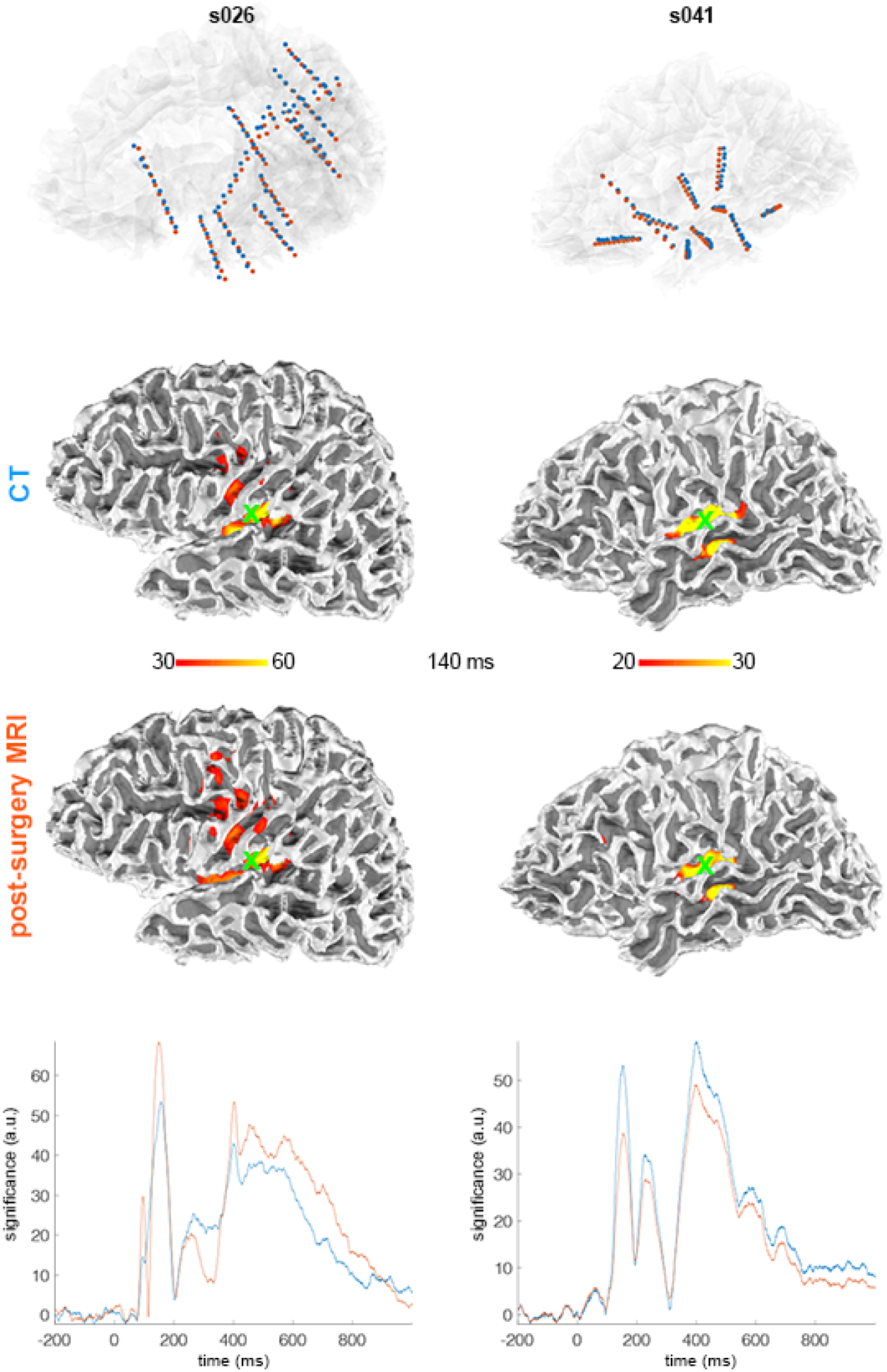
Source modeling based on the electrode contacts suggested by the post-surgery MRI and CT. Locations of the electrode contacts suggested by the post-surgery MRI and CT (top panel), the estimated neural current distributions using the electrode contact locations informed by CT (the second row from the top) and post-surgery MRI (the third row from the top), and the time courses of statistical significance in the auditory cortex from two representative patients.

## Discussion

Here, we describe a novel procedure to estimate a neural current distribution based on the discrete sampling of sEEG data with realistic anatomical information from MRI. In our empirical data with auditory stimulation, the estimated sources of activations were localized in the vicinity of auditory cortices of the superior temporal plane **(Figures 1** and **2)**. Notably, as we hypothesized, the source localization results were anatomically highly consistent across participants **(Figure 1**), even though the locations of electrode contacts were highly variable across individuals. Further, our source estimation conducted in the frequency domain revealed significantly increased 60-140 Hz HBBG in STG, and HBBG decreases in IFG, consistent with the analyses of the directly recorded sEEG signal **(Figure 3)**. These results suggest that distributed source modeling of sEEG data offers a powerful way to conduct anatomically normalized human neurophysiological research at the group level, and thus opens entirely new possibilities for basic and clinical neuroscience research with sEEG recordings.

### Sensitivity and SNR

In this study, we systematically examined the sensitivity and SNR distributions of the estimated neural current accounting for the recorded sEEG data. The results show that the sensitivity and the SNR of the sEEG recording depend strongly on the distance between the estimated area vs. electrode contact location **(Figures 4, 5**, and **6)**. When we considered individual electrode locations, the sensitivity and SNR can become tens or hundreds of times smaller if the current source locations in the model is as little as one millimeter away from the nearest electrode contact **(Figures 7** and **8)**. The sharp drop of SNR and sensitivity for a moderate increase in the minimal distance, together with the cross-validation analysis **(Table 1**), suggested that precise electrode implantation is important to record the desired neural activity at the target brain locations. However, as the neural currents were estimated from an ensemble of sEEG electrodes, the dependency of the sensitivity and SNR on the distances between the brain region of interest and all electrode contacts was more complicated: The sensitivity and SNR can be comparable when the median distance from a brain location to all electrode contacts differed by a few centimeters. Therefore, combining data from multiple sEEG electrode contacts greatly improves the sensitivity and SNR.

### Inclusion of deep brain source locations

Facilitated by the computational anatomy, we were able to examine the sensitivity and SNR distribution at regions with automatically segmented anatomical labels. We were particularly interested in the deep brain areas, where non-invasive MEG and EEG measurements have a very limited sensitivity ^23^. Consistent with the fact that sEEG electrode contacts are in the proximity of deep brain areas, we found that the sensitivity and the SNR in, for example, the thalamus and brain stem were comparable to those of several cortical locations **(Figure 6)**. This result supports the feasibility of using distributed source modeling of sEEG data to examine the interactions between cortical and subcortical areas.

We found significant neural currents in deep brain areas. However, we were not able to pinpoint the locations toward the medial geniculate body or inferior colliculus, part of the known sub-cortical areas involved in the auditory processing pathway. To better identify neural currents at these potential regions, we may thus need to limit the source space to the candidate locations as a prior constraint. This solution would, however, come with the price of neglecting neural currents from locations outside the modeled regions.

### Cross-validation

We cross-validated the accuracy of the source modeling of sEEG data using the leave-one-out approach. Note that this cross-validation is different from the “goodness-of-fit” in MEG/EEG analyses. In our cross-validation, the data used to estimate the source distribution and the data used for prediction were exclusive sets. On the contrary, the “goodness-of-fit” in MEG/EEG analyses was derived from the same set of measurement for both estimating the source distribution and predicting the measurement based on the estimated sources. Our crossvalidation suggested that about 30-50% of the variance can be explained by the proposed source modeling **(Table 1**). In other words, despite the limiting effects of the sharp drop-off of the sensitivity of sEEG at locations away from the recording sites **(Figures 7** and **8)**, the source modeling still explained a significant portion of the variance. This is probably because of the elicited neural activity is spatially distributed and the model favors a spatially extended source distribution to explain the measurements. In such a case, the L-2 norm-based distributed source modeling can reasonably describe the distribution of the neural activity at areas away from the implanted electrodes.

Potential factors related to the cross-validation performance reported here include the choice of the experimental paradigm, the forward model, and the source model. Here we used the neural responses to rather simple auditory stimuli (white noise bursts) with *a priori* well-known foci of response activity as the means to validate the predicted sEEG measurements. Such empirical validation is closely related to how much neural activity was actually elicited by the auditory stimuli. If the elicited neural activity is focal, it is difficult to estimate such an activity from measurements at other contact locations with minimal responses. Accordingly, if the elicited neural activity is spatially diffuse, the estimated the neural activity could have better matched the measurement at the left-out contacts. This potential confound should be considered in the interpretation of our results.

The forward model used in this study can mitigate the challenge of calculating the electric potential within the neural current volume ^24,25^. More complicated forward models, such as the one using the Finite Element Model^26^, or simpler models, such as the one assuming a simple spherical geometry and homogeneous conductivity ^14^, could provide a potential alternative for our choice for the trade-off between accuracy and computational complexity. Additional benefits in the forward calculations could be achieved by using recently introduced boundary element fast multipole methods, which simulate anatomically realistic head models with unprecedented numerical accuracy and speed ^27^. As for the source model, we used the MNE for its computational efficiency. Other choices, such as the one preferring focal neural current estimate by imposing a constraint to minimize its L-1 norm, such as the minimum-current estimate ^28^, may be further investigated in the future. Good performance in the crossvalidation analysis may only appear in experiments where spatially extensive neural currents are present. For experiments with a focal neural current distribution, such as that in the early response of sensory processing in normal subjects and inter-ictal discharges in epilepsy patients, the performance of cross-validation may be low, because estimating neural currents from contacts other than the one close to the focal neural current can be difficult. Yet again, in such cases, very strong *a priori* hypotheses of the probable source location are available, in contrast to the more complex cognitive processes.

### Registration

Previous studies using the combination of CT and MRI suggested that the accuracy of identifying the locations of electrode contacts was in the range of about 1 mm ^29^. In our study, we used the MRI before and after electrode implantation to identify the locations of sEEG electrodes. Specifically, focal dark spots in the post-surgery MRI were taken as the sites of electrode contacts. However, the distortion in the post-surgery MRI due to the susceptibility and the spatial resolution of MRI can confound the accuracy of the electrode contract localization. Using only MRI to inform the electrode contact location had the benefit of reducing the exposure to ionizing radiation in a CT scan. Without a gold standard, it is difficult to judge whether the electrode registration is more accurate with either MRI or CT. Both MRI and CT have distinct challenges in localizing electrode contacts: CT needs to be registered to MRI, and a smooth skull has limited features to warrant accurate registration between CT and MRI. The skull of MRI is an essential part may be difficult to be described accurately because of the concerns on MRI spatial resolution (about 1 mm) and distortion due to systematic (such as the nonlinearity property in MRI gradient coils ^30^) and physiological (such as the susceptibility discontinuity between the brain, cerebrospinal fluid, skull, and scalp interfaces) reasons ^31^.

### Source estimates of high-frequency broadband gamma activity

Our results suggest that distributed source estimation could also be utilized for analyses of HBBG activity. Evidence for significantly increased HBBG to sound stimuli was found in the vicinity of auditory cortices. Interestingly, we found evidence of decreased HBBG in IFG and nearby areas right after the onset of the auditory stimulus, an effect that could reflect suppression of involuntary attention networks due to the repetitive nature of the non-target auditory stimulus for reviews, see ^32,33^. The most important result of the present analysis, however, was the considerable consistency of the source estimates of HBBG and the electrode data obtained from contacts in the vicinity of STG and IFG. These novel results suggest that distributed source modeling analyses is a way to conduct anatomically-normalized group analyses of this highly essential neural activity, which is believed to constitute a direct correlate of local firing activity in the human brain ^16–18^.

### Future studies

There are a few issues worth studying further. For example, it is not known how the regularization parameter modulates the current source estimate. The regularization parameter is crucial in deriving a stable solution because of the ill-posed nature of the lead fields in extracranial MEG/EEG measurements. However, in sEEG, the lead fields can vary significantly across patients due to different electrode implantation scheme. Further, the conditioning number of the lead field matrix in sEEG can be less ill-posed. Thus, how to optimize the regularization parameter is still an open question. Second, we derived the group inference by averaging neural current estimates across patients. Given quantitative estimates of sensitivity and SNR, it might be reasonable to weight the neural current estimates by sensitivity or SNR. However, without a gold standard, it is still difficult to justify whether such SNR or sensitivity weighted group average is a better choice.

### Conclusions

While sEEG provides unsurpassed spatiotemporal accuracy, the interpretation of results has been complicated because the loci of electrode implantations differ greatly across individual patients. We developed a distributed, anatomically realistic sEEG source-modeling approach with which it becomes possible to estimate both iERP and HBBG responses in any given location at group level. High sensitivity and SNR values were found both in cortical and subcortical source estimates. After logarithmic transformations, the sensitivity and SNR were linearly inversely related to the minimal distance between the brain location and electrode contacts (slope≈-3.6). The HGGB source estimates were remarkably consistent with analyses of intracranial-contact data. Distributed sEEG source modeling of iERP and HBBG responses provides a new powerful neuroimaging tool that opens up a wealth of possibilities for both basic and clinical neuroscience research.

## Materials and Methods

### Participants

This study was approved by the Institute Review Board of National Yang Ming University and Taipei Veteran General Hospital. Ten medically refractory patients (age: 21 – 45; nine female) gave written informed consent before participating in this study. Two patients were excluded for the analysis because a large portion of the brain was resected in previous surgeries. The number of the implanted electrode contacts and the hemisphere with electrodes, together with the patient’s demographic information, are listed in **Table 2**.

**Table 2.** Demographic, medical, and surgical information for patients.

| Patient | Age | Sex | # of electrode contacts (*N*) | electrode hemisphere |  | Epilepsy |
| --- | --- | --- | --- | --- | --- | --- |
|  |  |  |  | left hemi. | right hemi. | diagnosis |
| s026 | 25 | M | 69 | + | - | Left TLE |
| s027 | 29 | F | 104 | + | - | Left TLE |
| s031 | 33 | F | 95 | - | + | Right temporal<br>PNH and right<br>frontal<br>opercular PMG |
| s032 | 23 | F | 67 | + | - | Left TLE |
| s033 | 21 | F | 76 | + | - | Left TLE |
| s034 | 27 | F | 100 | - | + | Right frontal<br>and temporal<br>schizencephaly |
| s036 | 45 | F | 59 | - | + | Right TLE |
| s041 | 39 | F | 70 | + | + | Right TLE |
| max. | 45 |  | 104 |  |  |  |
| min. | 21 |  | 59 |  |  |  |

**TLE: temporal lobe epilepsy, PNH: periventricular nodular heterotopia, PMG: polymicrogyria.**

### Experiment design

Two runs of data were collected from each patient. Each run lasted for six minutes. Within each run, fifty trials of auditory stimuli, including 45 trials of white noise (equal power between 20 Hz and 10,000 Hz; 0.3 s duration) and five trials of pure tone (440 Hz; 0.3 s duration), were randomly presented. The minimum and the average inter-stimulus intervals were 1.2 s and 2.0 s, respectively. Patients were instructed to press a button when hearing a pure tone while ignoring the white noise stimuli. In this study, we only calculated the responses evoked by the white noise in order to avoid confounds related to motor responses. Auditory stimuli were delivered by an earphone (Model S14, Sensimetrics, Gloucester, MA, USA) using E-Prime (Psychology Software Tools, Sharpsburg, PA, USA).

### sEEG recording

The placement of the electrode was solely based on the patient’s benefit in identifying epileptogenic zones. Each patient was implanted with 8 or 10 electrodes, whose contacts were mostly distributed between bilateral temporal lobes. Each electrode (0.3 mm diameter and spacing between contact centers 5 mm; Ad-Tech, Racine, WI, USA) had 6 or 8 contacts. sEEG data were sampled at 2,048 Hz with an electrode at FPz as the reference.

### MRI acquisitions

*T*_1_-weighted MRI was collected before and after the surgery for electrode implantation on 3T MRI scanners (Skyra, SIEMENS, Erlangen, Germany; Discovery MR750, General Electric, Milwuakee, WI, USA). The imaging parameters were the same in two acquisitions: MPRAGE sequence, TR/TI/TE/flip = 2530 ms/1100 ms/3.49 ms/7°, partition thickness = 1.33 mm, matrix = 256 x 256, 128 partitions, FOV = 21 cm x 21 cm. Two sets of MRIs were acquired for each patient before and after electrode implantation.

### CT acquisitions

Two patients were acquired with CT after electrode implantation. CT images were used to guide the identification of electrode contact locations using 3D slicer^34^. Parameters used in the CT acquisition were 64 slices, rotation duration of 1 second with coverage of 16 cm per rotation, 60-kW generator (512 × 512 matrix), 120 kV, 301 mAs, and axial slice thickness of 1 mm.

### Data analysis

The first step of our analysis was to identify the location of electrode contacts in the individual’s brain. Specifically, in the post-surgery MRI, there were discrete dark image voxel clusters caused by the susceptibility artifact for each electrode contact. These dark image voxels clusters were used to identify the locations of contacts. We developed an in-house software in *Matlab* (version 2019b, MathWorks, Natick, MA, USA) with a graphical user interface to facilitate this process. Specifically, after manually specifying the distance between neighboring contacts and the number of contacts on an electrode, an electrode was moved around the whole brain such that contact locations matched dark image voxel clusters in the post-surgery MRI. Upon completing the manual identification of electrode locations, contact locations were further optimized by allowing minor translation (±10 mm) and rotation (±2 degrees) by minimizing the sum of squares of image voxel values at all contact locations and their neighboring 26 image voxels within a 3-by-3-by-3 image voxel cubic in the post-surgery MRI using the *patternsearch* function in *MATLAB*.

Identified electrode contact locations were registered to pre-surgery MRI, which was used to build Boundary Element Models (BEM’s) required for the lead field calculation and to define locations of potential neural current sources. In the construction of BEM’s, the inner-skull, out-skull, and outer-scalp surfaces were automatically created by FreeSurfer (http://surfer.nmr.mgh.harvard.edu) based on the pre-surgery MRI. We did not use the postsurgery MRI for BEM construction because of the concern on the susceptibility artifact caused by electrode contacts and surgery wounds. The cortical source space for each patient was created at the gray matter and white matter boundary with approximately 5-mm separation between neighboring source locations. In addition, we also had sub-cortical source space, including thalamus, caudate, putamen, hippocampus, amygdala, accumbens areas, and substancia nigra. These sub-cortical areas were automatically segmented from *T*_1_-weighted MRI ^35^. At each source location, we had three orthogonal neural current dipoles in +x, +y, and +z directions. The current source space, including both sources at cortical and sub-cortical areas, electrode contacts, and three surfaces of inner-skull, out-skull, and out-scalp from a representative patient are shown in **Figure 1 (A)**. With defined source space, electrode contact locations, and skull as well as scalp boundaries, the lead fields were calculated by the *OpenMEEG* package (https://openmeeg.github.io/)^24,25^.

The measured sEEG data and the neural current sources at time *t* were related to each other by

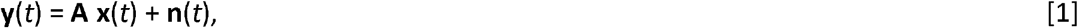

where **y**(*t*) denoted the collection of sEEG data across electrode contacts, **x**(*t*) denoted the neural current strength, and **n**(*t*) denoted the contaminating noise. In this study, we excluded the electrode contacts potentially related to epileptic activity when we created **y**(*t*). Note that **x**(*t*) had 3×*m* elements to describe the neural currents in three orthogonal directions at *m* brain locations. **A** was the lead field matrix. Specifically, for a unit current dipole source at location **r’** in the +*x*, +*y*, or +*z* direction, the electric potentials measured at all electrode contacts were

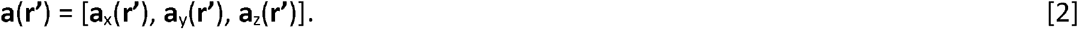

Assembling **a(r’)** across all possible current dipole source locations created the lead field matrix **A**.

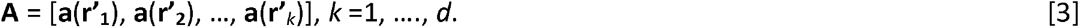

where *d* denotes the total number of current dipole source locations.

To estimate **x**(*t*) using the MNE, we had

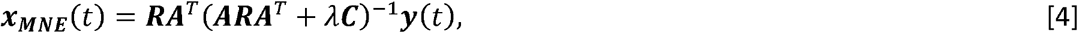

where **C** was the noise covariance matrix

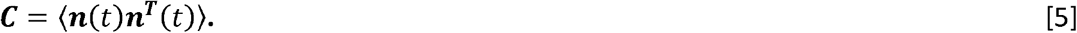

The operator 〈·〉 takes the ensemble average across realizations. In practice, **C** was estimated from **y**(*t*) during the pre-stimulus interval (from −200 ms to 0) with data concatenated across trials. The regularization Λ tuned the balance between the strength of the estimated neural current strength and the discrepancy between the modeled and measured data. We chose *λ* = 10 in this study as suggested by a previous MEG study ^13^.

The spatial distribution of estimated neural currents at each time instant from each patient was then spatially registered to an arbitrarily selected subject. Here we chose the subject “*fsaverage*” in the FreeSurfer library as the target subject. This registration was done via a spherical coordinate system ^36^. The neural currents were averaged across patients for each condition separately. The significance of neural current distribution was estimated by calculating the ratio between the instantaneous value and the standard deviation of the baseline interval at each source location after subtracting the mean of the estimates in the prestimulus interval. These ratios constituted the dynamic statistical parametric maps (dSPM) and were reported to follow a *t*-like distribution ^37^.

*We* quantified the spatiotemporal distribution of neural HBBG activity in the frequency domain. Specifically, after obtaining ***x_MNE_***(*t*) for each trial of stimulus, a Morlet wavelet function was applied to ***x_MNE_***(*t*) to extract frequency-specific HGGB signal at the central frequency *f*. The frequency selectivity was controlled by using five cycles of waveform. In this study, we specifically focused on the putative “non-oscillatory” HBBG between 60 Hz and 140 Hz. After the Morlet wavelet filtering, we took the absolute values of the frequency specific HBBG waveforms. At each brain location, we calculated the root of the sum of squares (RSS) of these wavelet-filtered waveforms of three directional components. Then, we averaged RSS of the frequency-specific wavelet-filtered waveforms across trials. The average waveform was then normalized to the average of the pre-stimulus baseline interval. We took the 10-based logarithm of the normalized average waveform as the spatiotemporal map of HBBG at frequency *f* for each patient. Similar to the group dSPM, these maps were combined across patients by averaging, subtracting the mean of the average, and dividing the standard deviation in the pre-stimulus baseline interval.

These calculations were validated by examining the correspondence of the HBBG between the source modeling from a group of patients and measurements from implanted electrodes. Specifically, we found patients with electrodes implanted at both the left superior temporal gyrus (STG) and inferior frontal gyrus (IFG). The HBBG was estimated by the source modeling of measurements from all patients except the chosen one. Then, HBBG from the dSPM were compared with the results from electrodes close to the left STG and IFG.

### Sensitivity and signal-to-noise ratio evaluation

With a given distribution of implanted electrodes and a set of current dipole source locations, we defined the sensitivity for a current dipole source at location **r’** as

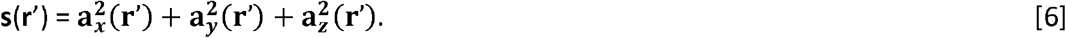

A map of **s(r’)** across current dipole source locations quantitatively depicted locations in the brain most sensitive to a particular set of implanted electrodes.

The signal-to-noise (SNR) at a specific location in the brain was derived from the “whitened” lead field matrix **A**. We used Singular Value Decomposition on the noise covariance matrix **C** to obtain a whitening matrix, which was then used to remove the correlation among lead fields and to normalize the sensitivity.

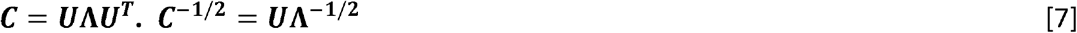

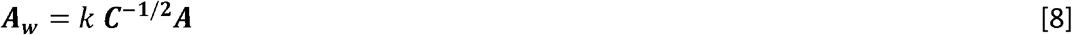

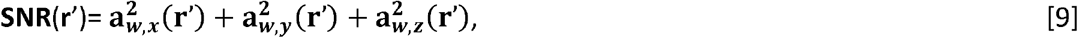

where **a_*w,x*_(r’)**, **a_*w,y*_(r’)**, and **a_*w,z*_(r’)** were columns of ***A_w_*** corresponding to the location **r’** with lead fields in the *x*-, *y*-, and, *z*-direction, respectively. *k* denotes a scaling factor. Because the lead field matrix ***A*** was calculated based on the theory and the noise covariance ***C*** was derived from empirical data, there was no information regarding the relative contribution from both terms. Thus, we arbitrarily chose *k* =1 in this study. The interpretation of the SNR should be careful considering the possible strength of the neural current and the noise level in different measurement conditions.

### Cross-validation

We used cross-validation to evaluate the accuracy of source modeling. Specifically, in a patient, the sEEG data at one contact were left-out, whereas a part of the remaining contacts was used for source modeling. The estimated sources were then used to predict the measurement at the left-out electrode contact by multiplying its lead field matrix and the estimated neural current distribution. Such synthetic data were compared with the actual measurement at the left-out contact. Each contact was taken as the left-out contact in each analysis, where the included contacts for source modeling were randomly selected and parametrically varied between 100%, 90%, 70%, and 50% of the remaining contacts. Ideally, the measurement at the left-out contact and the synthetic data should match each other. We used the percentage mean-squared-error as a metric to evaluate how much information was lost or retained in the source modeling.

### Stability of the source modeling

The locations of electrode contacts were identified by the guidance of post-surgery MRI, where focal black spots were taken as the locations of electrode contacts because of the susceptibility effects. This procedure was recommended in a previous study ^7^ and used to all patients in this study. Alternatively, electrode contacts were localized by the guidance of computer tomographic images after registered, with the help of the FSL (*flirt* function) package, and fused with the pre-surgery MRI. Two patients also followed this procedure to localize electrode contacts. We measured the average, standard deviation, minimal distance, and maximal distance between the discrepancy between two sets of electrode contact locations. Two different lead field matrices were calculated accordingly. We calculated the neural currents using these two lead field matrices and compared the differences in the estimated spatiotemporal dynamics of neural activity.

## Acknowledgments

This study was supported by Ministry of Science and Technology, Taiwan (103-2628-B-002-002-MY3, 105-2221-E-002-104), National Health Research Institutes (NHRI-EX108-10727EI), the Academy of Finland (project numbers 138145 and 276643 to IPJ and 298131 to FHL), aivoAALTO project funding of the Aalto University, National Institute on Deafness and other Communication Disorders (R01DC016765 supported JA, FHL, WJK; R01DC017991 supported JA, IPJ; R01DC016915 supported JA) and Russian Federation Government grant ag. 075-15-2019-1930 to I.P.J.

## References

1 Buzsaki, G., Anastassiou, C. A. & Koch, C. The origin of extracellular fields and currents--EEG, ECoG, LFP and spikes. Nat Rev Neurosci 13, 407–420, doi:10.1038/nrn3241nrn3241 [pii] (2012).

2 Bartolomei, F. et al. Defining epileptogenic networks: Contribution of SEEG and signal analysis. Epilepsia 58, 1131–1147, doi:10.1111/epi.13791 (2017).

3 Serletis, D., Bulacio, J., Bingaman, W., Najm, I. & Gonzalez-Martinez, J. The stereotactic approach for mapping epileptic networks: a prospective study of 200 patients. J Neurosurg 121, 1239–1246, doi:10.3171/2014.7.JNS132306 (2014).

4 Piai, V. et al. Direct brain recordings reveal hippocampal rhythm underpinnings of language processing. Proc Natl Acad Sci U S A 113, 11366–11371, doi:10.1073/pnas.1603312113 (2016).

5 Tang, C., Hamilton, L. S. & Chang, E. F. Intonational speech prosody encoding in the human auditory cortex. Science 357, 797–801, doi:10.1126/science.aam8577 (2017).

6 Zheng, J. et al. Amygdala-hippocampal dynamics during salient information processing. Nat Commun 8, 14413, doi:10.1038/ncomms14413 (2017).

7 Stolk, A. et al. Integrated analysis of anatomical and electrophysiological human intracranial data. Nat Protoc 13, 1699–1723, doi:10.1038/s41596-018-0009-6 (2018).

8 Gonzalez-Martinez, J. et al. Stereotactic placement of depth electrodes in medically intractable epilepsy. J Neurosurg 120, 639–644, doi:10.3171/2013.11.JNS13635 (2014).

9 De Momi, E. et al. Automatic trajectory planner for StereoElectroEncephaloGraphy procedures: a retrospective study. IEEE Trans Biomed Eng 60, 986–993, doi:10.1109/TBME.2012.2231681 (2013).

10 Cardinale, F. et al. Stereoelectroencephalography: surgical methodology, safety, and stereotactic application accuracy in 500 procedures. Neurosurgery 72, 353–366; discussion 366, doi:10.1227/NEU.0b013e31827d1161 (2013).

11 Hamalainen, M. S. & Ilmoniemi, R. J. Interpreting magnetic fields of the brain: minimum norm estimates. Med Biol Eng Comput 32, 35–42, doi:10.1007/bf02512476 (1994).

12 Goldenholz, D. M. et al. Mapping the signal-to-noise-ratios of cortical sources in magnetoencephalography and electroencephalography. Hum Brain Mapp 30, 1077–1086, doi:10.1002/hbm.20571 (2009).

13 Lin, F. H., Belliveau, J. W., Dale, A. M. & Hämäläinen, M. S. Distributed current estimates using cortical orientation constraints. Hum Brain Mapp 27, 1–13 (2006).

14 Yvert, B., Fischer, C., Bertrand, O. & Pernier, J. Localization of human supratemporal auditory areas from intracerebral auditory evoked potentials using distributed source models. Neuroimage 28, 140–153, doi:10.1016/j.neuroimage.2005.05.056 (2005).

15 Hosseini, S. A. H., Sohrabpour, A. & He, B. Electromagnetic source imaging using simultaneous scalp EEG and intracranial EEG: An emerging tool for interacting with pathological brain networks. Clin Neurophysiol 129, 168–187, doi:10.1016/j.clinph.2017.10.027 (2018).

16 Manning, J. R., Jacobs, J., Fried, I. & Kahana, M. J. Broadband shifts in local field potential power spectra are correlated with single-neuron spiking in humans. J Neurosci 29, 13613–13620, doi:10.1523/JNEUROSCI.2041-09.2009 29/43/13613 [pii] (2009).

17 Miller, K. J. Broadband spectral change: evidence for a macroscale correlate of population firing rate? J Neurosci 30, 6477–6479, doi:10.1523/JNEUROSCI.6401-09.2010 30/19/6477 [pii] (2010).

18 Ray, S. & Maunsell, J. H. Different origins of gamma rhythm and high-gamma activity in macaque visual cortex. PLoS Biol 9, e1000610, doi:10.1371/journal.pbio.1000610 (2011).

19 Cervenka, M. C., Nagle, S. & Boatman-Reich, D. Cortical high-gamma responses in auditory processing. Am J Audiol 20, 171–180, doi:10.1044/1059-0889(2011/10-0036) 20/2/171 [pii] (2011).

20 Iivanainen, J., Zetter, R. & Parkkonen, L. Potential of on-scalp MEG: Robust detection of human visual gamma-band responses. Hum Brain Mapp 41, 150–161, doi:10.1002/hbm.24795 (2020).

21 Parvizi, J. & Kastner, S. Promises and limitations of human intracranial electroencephalography. Nat Neurosci 21, 474–483, doi:10.1038/s41593-018-0108-2 10.1038/s41593-018-0108-2 [pii] (2018).

22 Hämäläinen, M. S. & Ilmoniemi, R. J. Interpreting magnetic fields of the brain: minimum norm estimates. Helsinki University of Technology, Espoo TKK-F-A559 (1984).

23 Baillet, S., Mosher, J. C. & Leahy, R. M. Electromagnetic brain mapping. IEEE Signal Processing Magazine 18, 14–30 (2001).

24 Gramfort, A., Papadopoulo, T., Olivi, E. & Clerc, M. OpenMEEG: opensource software for quasistatic bioelectromagnetics. Biomed Eng Online 9, 45, doi:10.1186/1475-925X-9-45 (2010).

25 Kybic, J. et al. A common formalism for the integral formulations of the forward EEG problem. IEEE Trans Med Imaging 24, 12–28, doi:10.1109/tmi.2004.837363 (2005).

26 Wolters, C. H. et al. Influence of tissue conductivity anisotropy on EEG/MEG field and return current computation in a realistic head model: a simulation and visualization study using high-resolution finite element modeling. Neuroimage 30, 813–826, doi:10.1016/j.neuroimage.2005.10.014 (2006).

27 Makarov, S. et al. Boundary Element Fast Multipole Method for Enhanced Modeling of Neurophysiological Recordings. IEEE Transactions on Biomedical Engineering, 1–1, doi:10.1109/TBME.2020.2999271 (2020).

28 Uutela, K., Hamalainen, M. & Somersalo, E. Visualization of magnetoencephalographic data using minimum current estimates. Neuroimage 10, 173–180, doi:10.1006/nimg.1999.0454 (1999).

29 Blenkmann, A. O. et al. iElectrodes: A Comprehensive Open-Source Toolbox for Depth and Subdural Grid Electrode Localization. Front Neuroinform 11, 14, doi:10.3389/fninf.2017.00014 (2017).

30 O’Donnell, M. & Edelstein, W. A. NMR imaging in the presence of magnetic field inhomogeneities and gradient field nonlinearities. Med Phys 12, 20–26, doi:10.1118/1.595732 (1985).

31 van der Kouwe, A. J. W., Benner, T., Salat, D. H. & Fischl, B. Brain morphometry with multiecho MPRAGE. Neuroimage 40, 559–569, doi:10.1016/j.neuroimage.2007.12.025 (2008).

32 Jääskeläinen, I. P., Ahveninen, J., Belliveau, J. W., Raij, T. & Sams, M. Short-term plasticity in auditory cognition. Trends Neurosci 30, 653–661 (2007).

33 Jääskeläinen, I. P. et al. Short-term plasticity as a neural mechanism supporting memory and attentional functions. Brain Res 1422, 66–81, doi:S0006-8993(11)01734-3 [pii] 10.1016/j.brainres.2011.09.031 (2011).

34 Narizzano, M. et al. SEEG assistant: a 3DSlicer extension to support epilepsy surgery. BMC Bioinformatics 18, 124, doi:10.1186/s12859-017-1545-8 (2017).

35 Fischl, B. et al. Whole brain segmentation: automated labeling of neuroanatomical structures in the human brain. Neuron 33, 341–355, doi:10.1016/s0896-6273(02)00569-x (2002).

36 Fischl, B., Sereno, M. I., Tootell, R. B. & Dale, A. M. High-resolution intersubject averaging and a coordinate system for the cortical surface. Hum Brain Mapp 8, 272–284, doi:10.1002/(sici)1097-0193(1999)8:4<272::aid-hbm10>3.0.co;2-4 (1999).

37 Dale, A. M. et al. Dynamic statistical parametric mapping: combining fMRI and MEG for high-resolution imaging of cortical activity. Neuron 26, 55–67, doi:10.1016/s0896-6273(00)81138-1 (2000).

